# Disentangling enabler and adaptive traits across habitat shifts in a large neotropical plant clade

**DOI:** 10.64898/2026.09.17.752501

**Authors:** Yacov Kilsztajn, Thais Vasconcelos, James D. Boyko, Vanessa Graziele Staggemeier

**Affiliations:** Programa de Pós-Graduação em Ecologia, Universidade Federal do Rio Grande do Norte, Natal/RN, Brasil; Department of Ecology and Evolutionary Biology, University of Michigan, USA; University of Michigan Herbarium, University of Michigan, USA; Departamento de Ecologia, Universidade Federal do Rio Grande do Norte, Natal/RN, Brasil

**Keywords:** biome-shifts, climatic niche, Myrteae, Myrtaceae, correlated trait evolution

## Abstract

- Plant lineages tend to retain their ancestral environments, yet some successfully shift into new habitats. However, the factors that enable certain lineages to colonize novel environments while others fail to do so remain unclear. We tested whether vegetative traits function as enablers of habitat transitions, whereas reproductive traits evolve primarily as adaptive responses following these shifts.
- We analyzed 12 vegetative and 7 reproductive traits across 689 species of Neotropical Myrtaceae. Habitats were characterized using biome categories and multiple climatic gradients. Using a set of trait evolution models, we quantified trait–environment matching and evaluated whether trait evolution preceded or followed habitat transitions.
- Vegetative traits showed stronger trait-environment matching and more frequently acted as enablers, while reproductive traits displayed both enabler and adaptive patterns. Transitions into harsher environments were generally associated with a greater number of enabler traits.
- Habitat shifts are associated with a combination of enabler and adaptive patterns that vary across trait types and environments. Reproductive traits may contribute to colonization more than often assumed, and colonization of harsh environments may require more enabler trait combinations, highlighting that different trait-evolutionary dynamics may underlie transitions into different habitats.

## Introduction

Environmental change plays a central role in shaping the evolutionary history and geographic distribution of plant lineages (Davis & Shaw, 2001; Hewitt, 2000; Ackerly, 2003; Donoghue, 2008). When environmental conditions shift through time, species may respond by tracking suitable habitats across space or by evolving the capacity to tolerate new ecological conditions (Donoghue, 2008; Weeks et al., 2014). The tendency of lineages to retain ancestral ecological tolerances (known as niche conservatism) has been widely documented, showing that many clades track particular environments over long evolutionary timescales (Wiens & Donoghue 2004; Wiens & Graham 2005; Losos 2008; Wiens et al. 2010; Crisp & Cook 2012). However, many plant lineages have successfully colonized new habitats through time, indicating that ecological tolerances are not always static (Donoghue, 2008; Donoghue & Edwards, 2014). This raises a central question in evolutionary ecology: what determines whether lineages remain confined to their ancestral environments or successfully shift into new habitats?

One factor proposed to influence the probability of such transitions is the presence of enabler traits (Donoghue 2005; Donoghue & Edwards, 2014). These traits may provide an initial level of tolerance that allows populations to persist long enough for natural selection to subsequently fine-tune other traits that optimize performance under the new selective regime, hereafter referred to as adaptive traits. Both enabler and adaptive traits should contribute to the frequent observation of trait-environment matching across habitats; that is, the tendency for certain traits to occur more frequently in particular environments across independent lineages. The concept of enabler traits is what once would have been called “pre-adaptations”, a designation rejected by Gould and Vrba (1982) due to its misleading implications, yet still prevalent in contemporary plant macroevolutionary literature (e.g., Velasquez-Puentes et al., 2023; Li et al., 2025). Here we adopt the enabler trait concept (*sensu* Donoghue, 2005) instead of “pre-adaptation” terminology, to emphasize the role of traits in facilitating habitat transitions rather than the specific selective pressures that initially drove the trait’s origin (for further discussion see Gould & Vrba, 1982; Donoghue, 2005). Nevertheless, we acknowledge that studies investigating enabler traits address essentially the same evolutionary patterns as those searching for “pre-adaptations” in the broader sense employed in recent literature (e.g., Velasquez-Puentes et al., 2023; Li et al., 2025).

Several empirical examples illustrate how enabler traits facilitate habitat transitions. Sclerophyllous leaves, characterized by high tissue density and thick cuticles, enabled several plant lineages to colonize Mediterranean climates (Verdú et al., 2003; Ackerly, 2004; Salvo et al. 2010). Similarly, underground storage organs and resprouting ability predisposed lineages to colonize fire-prone biomes (Simon et al., 2009; Keeley et al., 2011), while herbaceousness and small xylem conduits enabled transitions into freezing environments (Zanne et al., 2014). Deciduousness enabled shifts from tropical to temperate environments in Cannabaceae (Li et al., 2025), just as small leaflet size acted as an enabler for the colonization of arid regions in *Swartzia* (Velasquez-Puentes et al., 2023).

Different traits may play distinct roles during habitat shifts (e.g., Zanne et al., 2014; Velasquez-Puentes et al., 2023). Vegetative traits directly influence resource acquisition and stress tolerance (Wright et al., 2004; Díaz et al., 2016), and therefore are likely to affect whether a lineage can successfully establish in a new environment (e.g., Ackerly, 2004; Verdú et al., 2003; Simon et al., 2009; Salvo et al., 2010; Keeley et al., 2011; Zanne et al., 2014; Li et al., 2025). In contrast, reproductive traits often mediate interactions with pollinators and seed dispersers (Harder & Johnson, 2009; Rojas et al., 2022), and may evolve after habitat transitions as species adapt to novel ecological interactions present in the new environment (e.g., Hamilton & Wessinger, 2022; Velasquez-Puentes et al., 2023; Kopper et al., 2024; Dale et al., 2026).

Consequently, vegetative traits may more often act as enablers of habitat shifts, whereas reproductive traits may more frequently evolve as adaptive responses following environmental transitions. Although this reasoning is conceptually appealing, no study to our knowledge has explicitly tested it. Most studies investigating the timing of trait evolution relative to habitat shifts focus on a small number of traits, limiting this type of comparison (e.g. Ackerly, 2004; Verdú et al., 2003; Simon et al., 2009; Salvo et al., 2010; Keeley et al., 2011; Zanne et al., 2014; Hamilton & Wessinger, 2022; Velasquez-Puentes et al., 2023; Kopper et al., 2024; Li et al., 2025). Considering multiple traits, including many vegetative and reproductive traits, under the same framework may provide a more comprehensive understanding of how traits influence lineages’ transitions into new habitats.

To distinguish between enabler and adaptive traits across vegetative and reproductive structures, we used the Neotropical clade (*sensu* Vasconcelos et al., 2017) of the tribe Myrteae (Myrtaceae) as a model system (hereafter Neotropical Myrtaceae). This lineage encompasses nearly all Myrtaceae diversity in the Neotropics and includes approximately 2,500 species (NMWG et al., 2024). It occupies a wide range of habitats, from rainforests to seasonally dry environments (Oliveira-Filho & Fontes, 2000), and its species exhibit substantial variation in both vegetative and reproductive traits (Santos-Neves et al., 2023; Cunha, 2024). This morphological and ecological diversity makes Neotropical Myrtaceae an ideal system to investigate the relationship between habitat shifts and trait evolution, while also enabling broader inferences applicable to other plant clades, particularly tropical woody taxa.

Here, we investigate habitat shifts in the evolutionary history of Neotropical Myrtaceae to test whether the evolution of specific traits precedes or follows colonization of new environments. Specifically, we hypothesize that (H1) vegetative traits act as enablers of habitat shifts, whereas (H2) reproductive traits function as adaptive features evolving after environmental transitions.

## Material and Methods

### Sampling trait data

Considering the evolutionary scope of our study, we selected species based on their presence in phylogenetic hypotheses. Our sampling included 689 Neotropical Myrtaceae species (∼28% of the clade diversity) included in the most comprehensive phylogeny for Neotropical Myrtaceae available to date (NMWG et al., 2024). We focused on vegetative and reproductive morphological traits expected to be associated with environmental variation; i.e. to present trait-environment matching (Table S1). In total, 12 vegetative and 7 reproductive traits were measured based on 2,631 specimens (Table S2) using standard trait measurement procedure (Perez-Harguindeguy et al., 2016). Vegetative traits included leaf area, length/width ratio, circularity index, leaf mass per area (LMA), petiole and internode length, secondary vein density, phyllotaxis, and presence of cataphyll, trichomes, and drip-tip (acuminate leaf apex; Wang et al., 2020). Reproductive traits included flower, fruit, and seed size metrics (length and width, hypanthium and petal), number of flowers per inflorescence, presence of a terminal flower, fruit color in the mature stage, and number of seeds per fruit. Fruit color was treated as a discrete trait, with species classified as conspicuous (orange, red, black) or inconspicuous (brown, gray, green, yellow) based on their expected chromatic contrast against mature green foliage.

Leaf and flower traits were measured directly from physical herbarium specimens, primarily identified by expert Myrtaceae taxonomists (visited collections were ALCB, CEPEC, ESA, IAN, MBM, MG, RB, SP, SPF, UB, and UFRN). Some leaf traits (e.g., leaf area) were derived from digital images of the same specimens. To streamline image processing, we developed a custom Python script using OpenCV for automatic trait extraction (see Supporting Information, Methods S1). All outputs were manually checked, and for approximately one-third of the specimens (to which segmentation failed), leaf traits were manually measured in ImageJ (Schneider et al., 2012). For each species, we calculated mean trait values using data from 3–5 herbarium specimens whenever available (Table S2). Fruit and seed data were obtained from Cunha (2024). To account for instances in which multiple measurements existed for a single structure (fruit and seed), we estimated the volume using the ellipsoid formula: *(4/3)πd^2^l*, where *d* and *l* represent the width and length radii, respectively. For floral traits, we used a Principal Component Analysis (PCA) to summarize size variation based on petal length as well as the width and length of both the flower and the hypanthium. The first principal component (PC1) from the PCA explained 86.69% of the total variance and was used in subsequent analyses. All floral size traits loaded positively and similarly on PC1 (loadings ∼ 0.45), indicating that higher PC1 scores correspond to larger flowers (Fig. S1).

For species not represented in the visited herbaria, we extracted measurements from digitized herbarium sheets available through Global Biodiversity Information Facility (GBIF) and SpeciesLink (GBIF, 2025; SpeciesLink, 2025). However, as leaf weight could not be obtained from these images, we were unable to calculate LMA directly for 188 species (27% of all sampled species). To overcome this, we imputed missing LMA values using predictive models based on other leaf morphological traits (leaf area and petiole dimensions). We first evaluated a set of linear regression models used in paleobotanical and paleoclimatic studies (Royer et al., 2007), then refined them by adding interaction terms and phylogenetic information to enhance predictive performance (see Supporting Information, Fig. S2, Methods S2).

### Gathering environmental data

To obtain environmental data, we first gathered species occurrence records from GBIF (GBIF.org, 2025). To account for taxonomic changes, we retrieved known synonyms for each target species (POWO, 2024). When two or more taxa have previously been synonymized, but appeared in widely separated clades within the phylogeny (NMWG et al., 2024), they were treated as distinct taxonomic entities. Nomenclature was standardized prior to data download using the R package *taxize* (Chamberlain & Szöcs, 2013), which resolves species names against the GBIF backbone taxonomy. After downloading the occurrence records, we processed the data to improve accuracy following a pipeline adapted from Vasconcelos et al. (2023) using the R packages *raster*, *sp*, and *rgeos* (Pebesma & Bivand, 2005; Hijmans et al., 2015). We removed duplicate entries, records located at political centroids (within a 75 km buffer), and occurrences falling outside each species’ native distribution range as defined by the Plants of the World Online database (POWO, 2024). To reduce spatial sampling bias, we applied a quadrant thinning filter, retaining up to three randomly selected records per species within each 10-minute grid cell. Following this cleaning process, we manually reviewed species distributions to identify and correct potential errors, such as misassigned natural distribution range (verified using species descriptions) and centroid records associated with small islands. In such cases, verified records were retained. For species not represented on GBIF, we conducted targeted searches on SpeciesLink and scientific publications to obtain occurrence records (Vasconcelos et al., 2018; Flickinger et al., 2022; Gonzales & Gonzales, 2023). We also extracted coordinates from specimens used in trait sampling when they were not available on GBIF (see Table S2 for the final number of occurrence records per species).

To align our analysis with established research on trait evolution and habitat shifts, we first classified species by biome (e.g. Simon et al., 2009; Crisp et al., 2011; Schmerler et al., 2012; Zanne et al., 2014; Dale et al., 2024). This approach allows for a direct comparison between observed trait patterns and broad environmental transitions. We overlaid species’ cleaned occurrences on the World Wildlife Fund (WWF) map of terrestrial biomes (Olson et al., 2001) and then classified each species based on canopy structure by calculating the proportion of its records falling in closed-versus open-canopy biomes. In order to do that, we binarized the original 13 terrestrial biomes into those two categories. Occurrence points falling within biomes predominantly characterized by closed-canopy physiognomies (“Tropical & Subtropical Moist Broadleaf Forests”, “Tropical & Subtropical Coniferous Forests”, “Temperate Broadleaf & Mixed Forests”, “Temperate Conifer Forests”, and “Boreal Forests/Taiga”) were scored as closed-canopy biomes. In contrast, occurrences in biomes with generally low canopy cover (“Tropical & Subtropical Grasslands, Savannas & Shrubland”, “Tropical & Subtropical Dry Broadleaf Forests”, “Temperate Grasslands, Savannas & Shrublands”, “Flooded Grasslands & Savannas”, “Montane Grasslands & Shrublands”, “Tundra”, “Deserts & Xeric Shrublands”, “Mediterranean Forests, and Woodlands & Scrub”, “Mangroves”) were scored as open-canopy biomes.

While biome types classified by canopy structure provide a broad ecological context, climatic variables capture finer-scale environmental gradients. Environmental data were extracted from occurrence points using the *raster* and *geodata* R packages (Hijmans et al., 2015, 2024). We then calculated species-level means for each climatic variable across all cleaned records. Following previous work on plant trait–climate relationships (e.g. Wright et al., 2004, 2017; Šímová et al., 2018), we extracted variables representing major climatic axes, including mean annual temperature, diurnal temperature range, temperature seasonality, annual precipitation, precipitation seasonality, altitude, and aridity index (WorldClim; Fick & Hijmans, 2017; Zomer et al., 2022). To further capture interactions between energy and water availability, we also included temperature of the wettest quarter and precipitation of the warmest quarter (WorldClim; Fick & Hijmans, 2017). These variables were chosen because they represent key dimensions of environmental variation likely to influence trait morphology, particularly through effects of temperature, moisture availability, and their interaction.

### Trait-environment matching

The distinction between enabler and adaptive traits is contingent on the existence of a correlation between trait and environment. If a trait is not associated with environmental conditions, it is not meaningful to ask whether it functions as an enabler or as an adaptive feature. Therefore, before identifying enabler and adaptive traits, we first verified whether the selected traits (chosen based on their expected theoretical association with environmental conditions, Table S1) were indeed correlated with environmental variation in Neotropical Myrtaceae.

We tested relationships between each plant trait and environmental conditions using multiple approaches. For biome categories (closed- vs open), we applied phylogenetic ANOVA for continuous traits using the R package *phytools* (Revell, 2024), and performed Likelihood Ratio Tests (LRT) to compare dependent and independent evolutionary models for categorical traits using the R package *corHMM* (Boyko & Beaulieu, 2021). To test trait-environment matching we fitted phylogenetic generalized least squares (PGLS) models for continuous traits, and we used phylogenetic logistic regression (MPLE) models for categorical traits, both implemented in the R package *phylolm* (Ho et al., 2016). We standardized climatic variables to zero mean and unit variation and log-transformed trait values to meet model assumptions and facilitate interpretation. In cases in which models without log transformation showed better performance based on residual diagnostics, we retained those instead.

In total, we tested associations between 19 traits (12 vegetative and 7 reproductive) and biome category and climatic variables. The goal of these analyses was to detect significant trait-environment matching that could be used to evaluate our hypothesis regarding enabler and adaptive traits. To account for inflation of the Type I error rate due to multiple testing, p-values from the resulting tests were adjusted globally using the Benjamini –Hochberg false discovery rate (FDR) procedure, with an FDR threshold of 0.05. Although we recognize that the full set of results (including both significant and non-significant associations) is informative, here we focus on associations that remained significant after FDR correction, which were subsequently used to test our hypotheses.

### Identifying enabler and adaptive traits

For each trait–environment matching that was statistically significant (adjusted p < 0.05), we further tested whether traits were already present when lineages transitioned into a given habitat (enabler traits) or instead evolved after the transition (adaptive traits). To our knowledge, this distinction remains difficult to assess using continuous trait data, as there are no widely applicable methods that explicitly model direction-specific evolutionary transitions in a way that allows testing for asymmetric sequences of change. Recent developments such as generalized dynamic phylogenetic models (GDPMs; Ringen et al., 2026) provide a powerful and flexible framework for modeling the joint evolution of multiple continuous traits, including directed dependencies and feedback. However, although GDPMs can infer contingencies among traits (e.g. first trait, then environment), they do not currently allow explicit separation of directional shifts (e.g., increases versus decreases along environmental gradients). As a result, while GDPMs can indicate whether a trait generally acts as an enabler or responds adaptively, they cannot distinguish direction-specific scenarios. In practice, a trait may facilitate transitions toward drier conditions but not toward more humid environments, and failing to separate these directions could lead to misinterpretation of enabler versus adaptive effects.

In contrast, discrete character evolution models can readily be used to test directional changes in trait evolution (e.g., Zanne et al., 2014; Velasquez-Puentes et al., 2023; Li et al., 2025), and thus provide a feasible framework to evaluate our hypothesis. Additionally, our dataset includes categorical traits and biome classification, which further motivated the use of a unified discrete framework across all variables. We acknowledge that retaining traits in their continuous form would likely improve inference, and that discretization represents a limitation of our study. To test the sensitivity of this discretization, we categorized all continuous traits and climatic variables using three threshold values (40%, 50%, and 60%), and repeated all analyses across the nine possible combinations of trait and environmental thresholds.

For each trait–environment pair, we fitted discrete character evolution models using the corHMMDredge function (Boyko, 2026) in the R package corHMM (Boyko & Beaulieu, 2021). This approach automates the exploration of alternative transition-rate structures by iteratively constraining or removing parameters, while applying penalized likelihood to discourage overparameterization (Boyko, 2026). We used an L1 regularization penalty (λ = 1), which penalizes high transition rates and promotes sparse transition matrices. We explored models both with and without rate heterogeneity (max.rate.cat = 2) by incorporating a hidden Markov model (HMM) framework, thereby reducing the risk of detecting spurious correlations (Boyko & Beaulieu, 2013). Model exploration proceeded until further simplification no longer improved model fit, with termination occurring when successive models differed by more than two AIC units. Inference was based on the single best-supported model (i.e., lowest AIC), without model averaging, as the generated models are not associated with distinct a priori biological hypotheses (see Boyko, 2026 for details). The transition-rate matrix of the best-fitting model was then used to simulate 100 stochastic character histories conditional on the observed tip states (following Bollback, 2006). Indices to detect enabler and adaptive traits were calculated for each stochastic mapping, and we then estimated their mean values to support interpretation (Fig. 1).

**Figure 1.**
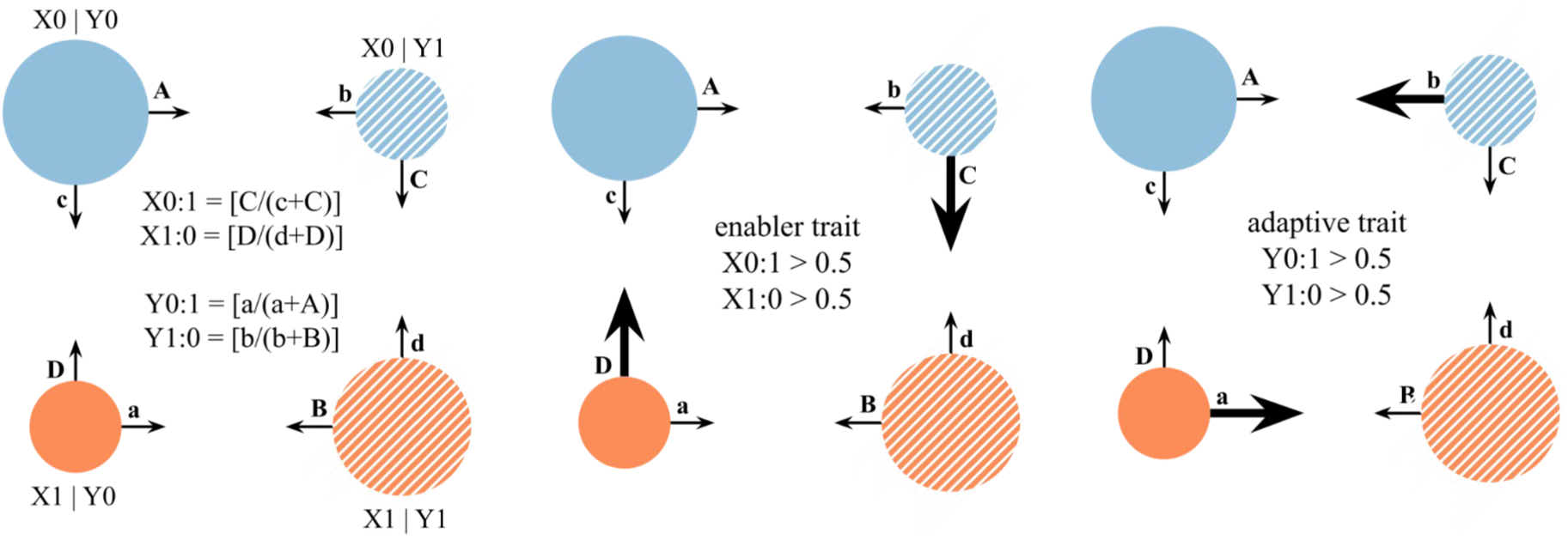
Theoretical framework for identifying enabler and adaptive traits based on transition count (a, b, c, d, A, B, C, D, E) from stochastic character mapping of discrete traits. Trait states are denoted as “Y0” (full circle) and “Y1” (dashed circle), and environmental states as “X0” (blue) and “X1” (orange). In this scenario the combinations X0Y0 and X1Y1 (larger circles) represent the most common — and presumably optimal — trait-environment associations, while thicker arrows denote the most frequent transitions.

To distinguish between enabler and adaptive traits, we adopted a decomposed framework that explicitly accounts for the directionality of both trait and habitat shifts. Rather than relying on a single general metric, we calculated transition-specific indices for each type of shift (for a detailed breakdown of the calculations, see Fig. 1). Environmental states were denoted as X0 and X1, and trait states as Y0 and Y1. Each index represents the proportion of a given transition relative to the total number of transitions of that type. Indices are named by combining X or Y (environment or trait, respectively) with the direction of change (e.g., 1:0 or 0:1), where the state before the colon denotes the ancestral condition and the state after the colon the derived condition.

Indices X1:0 and X0:1 quantify enabler effects relative to habitat shifts, whereas Y1:0 and Y0:1 quantify traits adaptive responses. Within this framework, and assuming that X0Y0 and X1Y1 represent the most frequent trait–environment associations, values greater than 0.5 indicate a predominance of enabler (for X1:0 and X0:1) or adaptive (for Y1:0 and Y0:1) patterns. All indices range from 0 to 1 and can be interpreted as the proportion of transitions consistent with either enabler (X indices) or adaptive (Y indices) process. For example, a value of X1:0 = 0.76 indicates that during transitions from environment X1 to X0, lineages already possessed the trait Y0 in 76% of cases, consistent with an enabler effect. More generally, environmental transition indices (X1:0 and X0:1) assess whether species already possess traits suited to a new environment prior to the shift (values > 0.5 indicating enabler), whereas intra-environmental transition indices (Y1:0 and Y0:1) evaluate whether trait evolution within a given environment is biased toward the most frequent (presumably optimal) state (values > 0.5 indicating potential adaptive evolution).

To facilitate the assessment of general patterns, we assigned binary states (“X0” or “X1”) to each environment based on their correlations with biome types (Fig. S3). We assigned “X1” to open-canopy biomes and environments characterized by low annual precipitation, temperature, and aridity index (i.e. dryer habitat), as well as high diurnal temperature range, altitude, precipitation and temperature seasonality (hereafter, xeric environments). Conversely, we assigned “X0” to closed-canopy biomes and environments with high annual precipitation, temperature, and aridity index (i.e. mesic habitat), and low diurnal temperature range, altitude, precipitation and temperature seasonality (hereafter, mesic environments). For each binarized trait, we assigned the state “Y0” to the trait value most frequent in “X0” environments, and “Y1” to the trait value most frequent in “Y1” environments. Thus, X0 represents mesic environments and Y0 the trait states most commonly associated with them, whereas X1 represents xeric environments and Y1 the trait states most commonly associated with those conditions. Consequently, X0:1 and X1:0 represent transitions from mesic to xeric environments and from xeric to mesic environments, respectively. Likewise, Y0:1 represents transitions toward trait states most commonly associated with xeric environments, whereas Y1:0 represents transitions toward trait states most commonly associated with mesic environments.

During the analysis of our results, we identified a potential bias in our indices. These indices assume that, under independent trait evolution, transitions between environment and trait states occur at equal rates regardless of state (X0:1, X1:0, Y0:1, Y1:0 = 0.5). However, even under independence, we noticed that transitions are more likely to occur in the most frequent state simply because it is more often available, making transition probabilities proportional to state frequencies (Fig. S4). This introduces a systematic bias, which becomes particularly relevant when varying thresholds alter trait state frequencies. To account for this, we implemented a null model that preserves the observed frequencies of trait and environmental states. Although the standard procedure would be to simulate data from an independent evolutionary model and then refit, such simulations do not guarantee recovery of the observed state frequencies, which was essential for our analyses.

For each trait–environment pair and its nine threshold combinations, we randomized tip states, explored alternative transition-rate structures using corHMMdredge, selected the best-supported model, and simulated character histories under that model. This procedure was repeated 100 times. Importantly, the best-supported model structure could differ among iterations, meaning that our null expectations incorporate the influence of state frequencies not only on parameter estimates, but also on model selection itself. We then compared observed indices to the null expectation by subtracting null values from observed values, yielding 100 differences per index for each threshold combination of each trait–environment pair. We further tested whether these differences deviated from zero using a Wilcoxon test. Given the large sample size, we additionally applied an effect size criterion: enabler or adaptive patterns were only inferred when the median difference was significant and at least 10 percentage points above the null expectation. Differences greater than 25 percentage points were classified as strong effects.

## Results

### Summary statistics

Vegetative and reproductive traits exhibited substantial interspecific variation. Leaf area ranged from 0.19 to 744.19 cm² (median: 12.14 cm²), while leaf mass per area (LMA) varied between 0.0062 and 0.0485 g/cm² (median: 0.0157 g/cm²). Petiole length ranged from 0.05 to 2.32 cm (median: 0.45 cm), and internode length from 0.23 to 13.48 cm (median: 2.23 cm). Leaf shape metrics were also highly variable: the circularity index ranged from 0.07 to 0.87 (median: 0.6), the length-to-width ratio from 0.9 to 19.34 (median: 2.3), and secondary vein density from 0.35 to 7.09 veins/cm (median: 1.73). Flower traits showed similarly broad variation. Petal length ranged from 0.68 to 21.89 mm (median: 2.79 mm), flower diameter from 2.36 to 49.73 mm (median: 8.00 mm), and flower length from 1.8 to 27.37 mm (median: 5.58 mm). Hypanthium diameter varied from 0.77 to 10.17 mm (median: 2.08 mm), and hypanthium length from 0.46 to 8.91 mm (median: 1.84 mm). The number of flowers per inflorescence ranged from 1 to 300 (median: 4 flowers). Fruit and seed traits also spanned a wide range. Fruit length ranged from 1.5 to 84.3 mm (median: 11.02 mm), fruit diameter from 2.5 to 62.0 mm (median: 10.31 mm), and fruit volume from 0.0932 to 763.52 cm³ (median: 3.24 cm³). Seed traits were similarly diverse. Seed length ranged from 1.38 to 40.0 mm (median: 6.50 mm), seed diameter from 0.75 to 38.09 mm (median: 5.81 mm), seed volume from 0.0021 to 130.45 cm³ (median: 0.52 cm³), and seed number per fruit from 1 to 264 (median: 2 seeds).

Categorical traits also exhibited marked variation across species. Trichomes were present in young leaves of 45.86% of species, while mature leaves retained them in 25.18%. Drip tips occurred in 46.98% of species, cataphylls in 40.52%, and opposite decussate phyllotaxis in 78.5%. Terminal flowers were observed in 36.47% of species, and 76.98% of species displayed conspicuous fruit colors (orange, red, or black).

### Trait-environment matching

In general, we found that several leaf, flower, fruit, and seed traits were associated with biome type (Fig. 2; Table S3), with vegetative traits showing a greater number of significant associations (7 of 12; 58%) than reproductive traits (2 of 7; 29%), after false discovery rate (FDR) adjustment. Species from closed-canopy biomes tend to have larger leaves with drip tips, longer petioles and internodes, opposite-distichous phyllotaxy and lower leaf mass per area (LMA), as well as more flowers per inflorescence and larger fruits. In contrast, species from open-canopy biomes typically exhibit smaller and thicker leaves without drip tips, shorter petioles and internodes, opposite-decussate phyllotaxy and higher LMA, fewer flowers per inflorescence, and smaller fruits.

**Figure 2.**
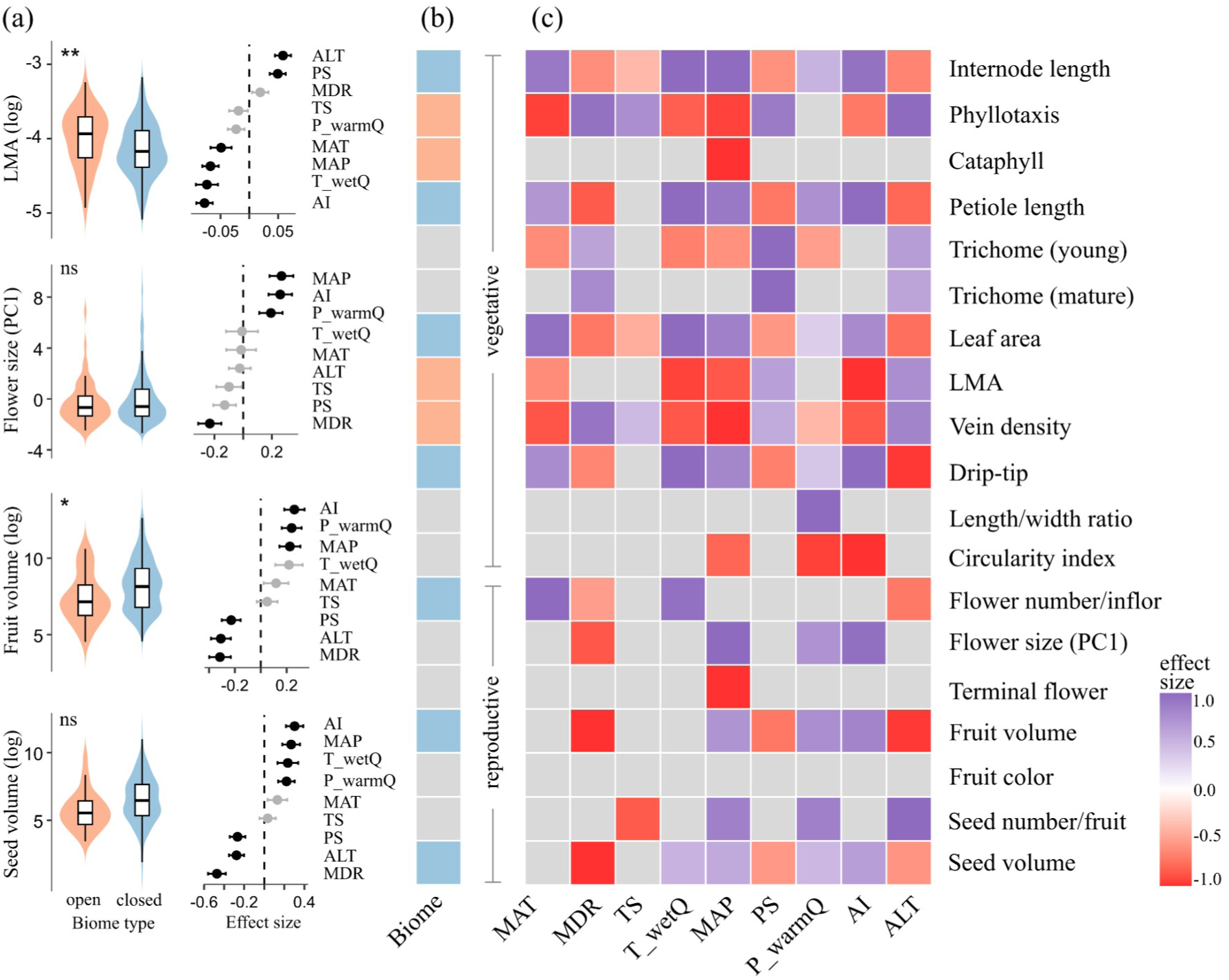
Trait–environment matching. Effect sizes correspond to beta coefficients from phylogenetic generalized least squares (PGLS) models. (a) Associations of leaf mass per area (LMA), flower size, fruit volume, and seed volume with biome type and climatic variables. Black dots indicate statistically significant effects (FDR-corrected p < 0.05). (b) Associations between all analyzed traits and biome type. Colors indicate the biome in which traits have higher values: closed canopy (blue; X0) or open canopy (orange; X1). Gray squares indicate non-significant associations (FDR-corrected p > 0.05). (c) Associations between all analyzed traits and climatic variables. Gray squares indicate non-significant associations (FDR-corrected p > 0.05). For discrete traits, positive effect sizes correspond to the following states: opposite-decussate phyllotaxy and presence of cataphylls, trichomes (in both young and mature leaves), drip tips, and terminal flowers. MAT: Mean Annual Temperature; MDR: Mean Diurnal Range; TS: Temperature Seasonality; T_wetQ: Temperature of the Wettest Quarter; MAP: Mean Annual Precipitation; PS: Precipitation Seasonality; P_warmQ: Precipitation of the Warmest Quarter; AI: Aridity Index; ALT: Altitude.

We detected consistent associations between plant traits and climatic variables across all plant organs (Fig. 2; Table S3), with vegetative traits again showing more significant relationships (72 of 102; 71%) than reproductive traits (25 of 63; 40%), after false discovery rate (FDR) adjustment. In environments characterized by high temperature and precipitation (i.e., more mesic habitats), species generally exhibit larger and narrower leaves with drip tips, lower LMA, and absence of trichomes in young leaves. These conditions are also associated with longer petioles and internodes, opposite-distichous phyllotaxy, lower secondary vein density, inflorescences lacking a terminal flower, more flowers per inflorescence, and larger flowers, fruits, and seeds, as well as a higher number of seeds per fruit. In contrast, habitats with high precipitation and temperature seasonality, greater diurnal temperature range, and higher altitude are associated with smaller leaves lacking drip tips and with higher LMA, as well as the presence of trichomes in both young and mature leaves. These environments also tend to be associated with opposite-decussate phyllotaxy, shorter petioles and internodes, higher secondary vein density, and smaller flowers, fruits, and seeds.

### Identifying enabler and adaptive traits

Across all significant trait-environment matchings (n = 106; see Table S3), patterns of enabler and adaptive effects varied with trait identity, environmental dimension, and the directionality of shifts (Fig. 3; Table S4). Transitions into open-canopy biomes were consistently associated with enabler traits reflecting reduced organ size and increased structural investment (e.g., small fruits, high LMA, and presence of cataphylls, and trichomes), whereas transitions to closed-canopy biomes were more frequent among lineages with large leaves and increased number of flowers per inflorescence (Fig. 3a). Adaptive responses following biome shifts were uncommon and were detected only for shifts toward smaller fruits in open-canopy environments (Fig. 3c).

**Figure 3.**
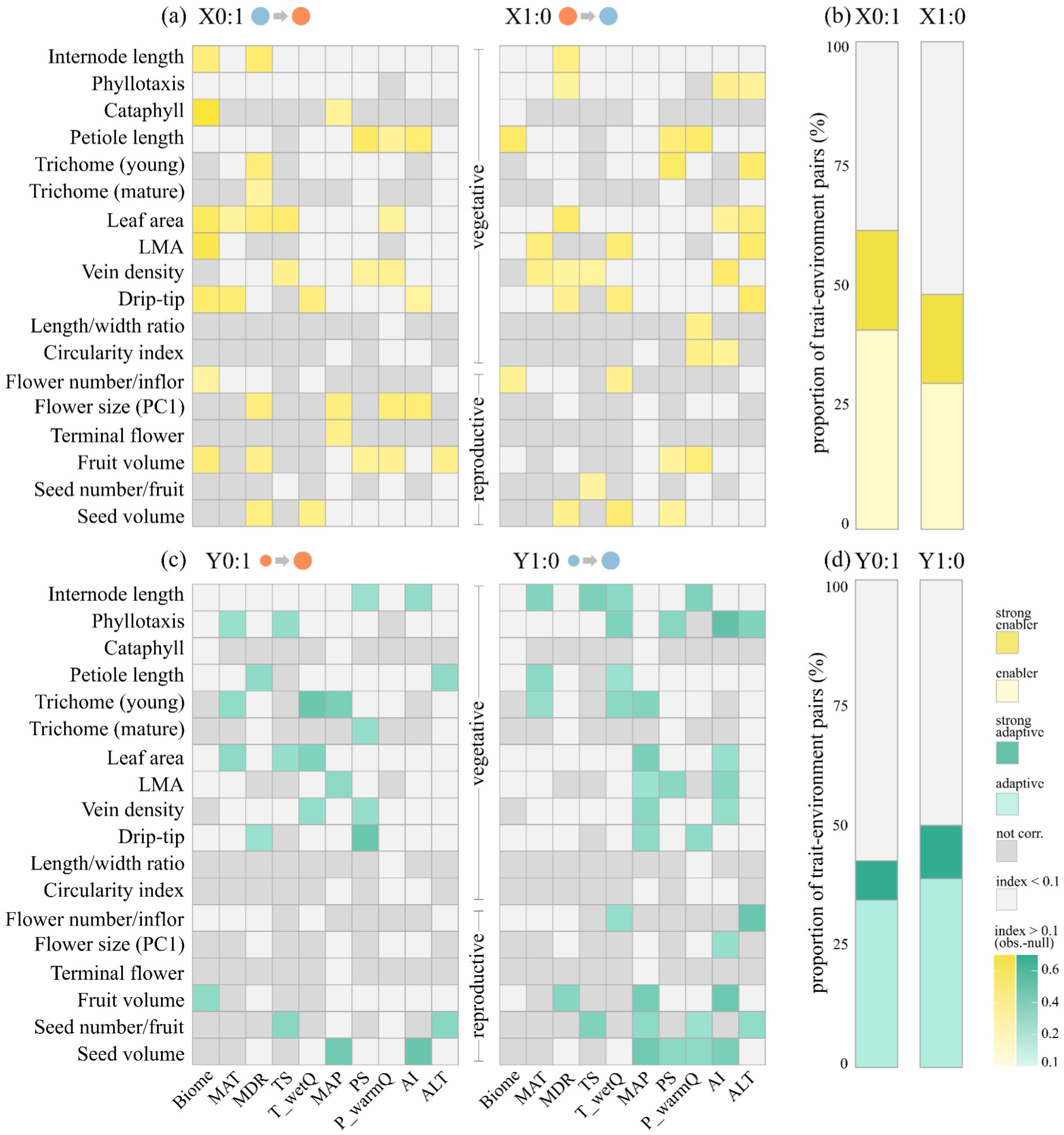
Differences between observed (obs.) and expected (null) transition indices for all analyzed traits, biome type, and climatic variables. (a) Median values across all threshold combinations for X0:1 (mesic to xeric) and X1:0 (xeric to mesic) indices used to identify enabler traits; (b) Proportion of trait–environment pairs exhibiting enabler (>0.1 above expected) and strong enabler (>0.25 above expected) effects; (c) Median values across all threshold combinations for Y0:1 and Y1:0 indices used to identify adaptive traits; (d) Proportion of trait–environment pairs exhibiting adaptive (>0.1 above expected) and strong adaptive (>0.25 above expected) effects. MAT: Mean Annual Temperature; MDR: Mean Diurnal Range; TS: Temperature Seasonality; T_wetQ: Temperature of the Wettest Quarter; MAP: Mean Annual Precipitation; PS: Precipitation Seasonality; P_warmQ: Precipitation of the Warmest Quarter; AI: Aridity Index; ALT: Altitude.

Across climatic gradients, transitions to dry environments were consistently enabled by traits associated with small organs (leaves, flowers, and fruits), and stress tolerance, including high LMA, presence of trichomes and cataphylls (Fig. 3a). In contrast, wet environments showed few enabler traits but strong adaptive responses toward acquisitive strategies, including large fruits and seeds, low LMA, low vein density, and presence of drip tips (Fig. 3c). Transitions to cold and seasonal environments were associated with enabler and adaptive traits reflecting size reduction and protection (e.g., high LMA, absence of drip tips, presence of trichomes, and smaller petioles), whereas warm and stable environments favored elongation (long internodes and petioles), and reduced structural investment (low LMA and vein density) (Fig. 3a,c).

Transitions toward xeric environments (X1; i.e., open-canopy biomes and conditions characterized by lower precipitation, as well as higher elevation and greater climatic seasonality) were associated with a greater number of enabler traits than transitions toward mesic environments (X0; i.e., closed-canopy biomes and conditions with higher precipitation, as well as lower elevation and reduced seasonality). Conversely, adaptive traits were more frequently detected in mesic than in xeric environments (Fig. 3 b,d).

Reproductive traits showed a higher frequency of enabler than adaptive patterns during transitions toward the optimal trait state in xeric environments (X1; Fig. 4a), however adaptive effects reached higher magnitudes than enabler effects (Fig. 4b). In contrast, for transitions toward the optimal trait state in mesic environments (X0), adaptive patterns were more frequent (Fig. 4a), and their magnitudes were comparable to those of enabler traits (Fig. 4b). Vegetative traits likewise showed a higher frequency of enabler than adaptive patterns during transitions toward the optimal trait state in xeric environments (Fig. 4a), with enabler effects also reaching higher magnitudes (Fig. 4b). For transitions toward the optimal state in mesic environments, differences in both frequency and magnitude between enabler and adaptive patterns were small, although strong enabler traits remained more common than strong adaptive ones (Fig. 4a).

**Figure 4.**
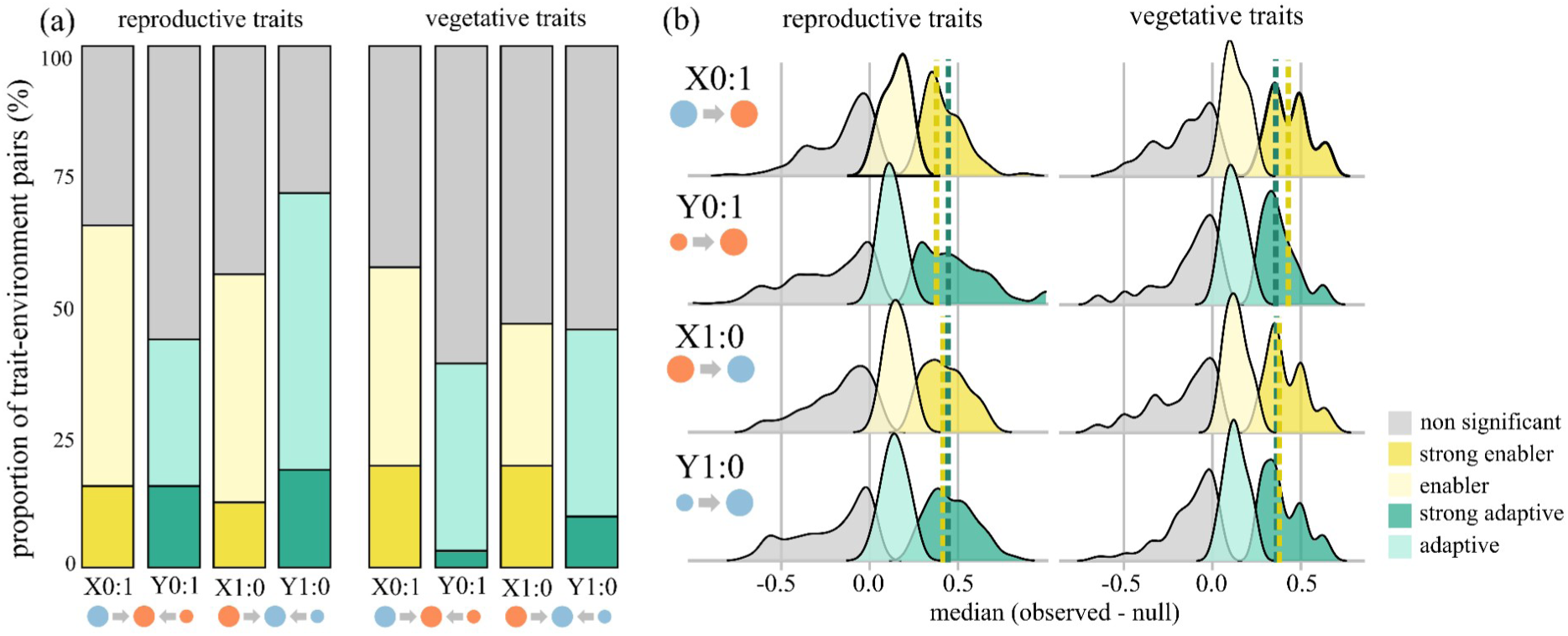
Differences between observed (obs.) and expected (null) transition indices for vegetative and reproductive traits. (a) Proportion of trait–environment pairs (median values across all threshold combinations) exhibiting enabler/adaptive (>0.1 above expected) and strong enabler/adaptive (>0.25 above expected) effects. (b) Median values (across all threshold combinations) of enabler/adaptive (>0.1 above expected) and strong enabler/adaptive (>0.25 above expected) effects for vegetative and reproductive traits; dashed lines represent median values for strong enabler/adaptive effects.

## Discussion

Our results partially support H1 (that vegetative traits act as enablers of habitat shifts) and H2 (that reproductive traits respond adaptively to habitat transitions). Vegetative traits more frequently exhibited enabler patterns and showed stronger enabler effects overall. In contrast, reproductive traits tended to show more adaptive patterns in shifts toward closed-canopy, more humid, warmer, and less seasonal habitats (environments coded as X0), but also displayed enabler effects, particularly during transitions to open-canopy, drier, colder, and more seasonal habitats (environments coded as X1). Additionally, we identified an unexpected pattern: transitions to xeric environments (X1) were more often associated with enabler traits and less frequently with adaptive traits compared to transitions to mesic environments (X0).

### Trait-environment matching is more often observed in vegetative traits

Vegetative traits were more consistently associated with the environment than reproductive traits. This pattern is expected because abiotic selective pressures are generally thought to act more strongly on vegetative structures than on reproductive ones (Berg, 1960; e.g., Dorken & Barrett, 2004; Mal & Lovett-Doust, 2005; Brock & Weinig, 2007; Pélabon et al., 2011). Reproductive structures such as flowers, fruits, and seeds are often subject to stronger biotic selective pressures relative to abiotic factors (e.g., Valenta et al., 2020; Basnet et al., 2025; Cabral et al., 2025), whereas vegetative traits, although also influenced by biotic interactions such as herbivory (e.g., Brown & Lawton, 1991; Hulshof et al., 2013; Turcotte et al., 2014), tend to be more directly shaped by abiotic environmental conditions (e.g., Poorter & Rozendaal, 2008; Meng et al., 2009; Sack et al., 2012; Li et al., 2015; Maracahipes et al., 2018; Han et al., 2019; Wang et al., 2020; Filartiga et al., 2022; Wang et al., 2022).

An additional notable pattern in our results is that functionally distinct traits often exhibited similar associations with environmental gradients. For example, internode and petiole length, as well as leaf area, showed highly similar responses to environmental variation. Likewise, fruit volume, and seed volume displayed closely aligned patterns across environmental variables. These coordinated responses may reflect developmental and functional integration among traits (Olson & Miller, 1958; Berg, 1960; Primack, 1987), whereby selection acting on one trait may indirectly influence others (Gianoli & Palacio-López, 2009). However, coordinated responses do not necessarily imply that traits contribute equally or simultaneously to habitat-shifts. Although integrated traits often evolved in the same direction, their apparent roles differed depending on the environmental context. For instance, smaller leaves and fruits appear to have facilitated transitions into environments with high precipitation seasonality, whereas petiole and internode length, as well as seed size, seem less influential in these shifts. In contrast, transitions into environments with high diurnal temperature range are more often associated with longer petioles and internodes, as well as larger fruits and seeds, while leaf area appears to play a more limited role. These patterns are consistent with the concept of mosaic evolution, which emphasizes that different traits need not evolve at the same pace or under the same selective pressures (Stebbins, 1984). Rather than evolving as fixed adaptive syndromes, integrated traits may exhibit similar long-term responses to environmental gradients while differing in the timing of their evolutionary change. Consequently, some traits may facilitate habitat-shifts by changing early, whereas others respond only after lineages have colonized new environments (Donoghue & Edwards, 2014).

It is important to note that our analyses did not explicitly model trait covariance. Approaches that incorporate trait integration may reveal additional structure in these relationships and provide a more comprehensive understanding of how traits jointly influence habitat transitions. Beyond identifying individual traits that facilitate habitat-shifts, investigating how the strength of trait integration evolves across lineages and influences the likelihood of habitat-shifts represents a promising avenue for future research (Parins-Fukuchi, 2020; Parins-Fukuchi et al., 2024).

### Reproductive traits also exhibit enabler effects

Although vegetative traits are often emphasized as key determinants in allowing plant lineages to arrive, survive and establish in certain habitats (e.g., Verdú et al., 2003; Ackerly, 2004; Simon et al., 2009; Salvo et al., 2010; Keeley et al., 2011; Zanne et al., 2014; Velasquez-Puentes et al., 2023; Li et al., 2025), our results suggest that reproductive traits may also frequently exhibit enabler patterns, particularly during transitions into open-canopy, drier, colder and more seasonal habitats. This differs from our initial expectations and indicates that successful habitat shifts may depend not only on the ability to establish and persist under new environmental conditions, but also on the capacity to reproduce effectively within them (Theoharides & Dukes, 2007; Barrett, 2011). If reproductive structures are not suited to the new habitat (e.g., due to high energetic costs or mismatches with pollinators or seed dispersers) species may establish and grow vegetatively but still experience reduced reproductive success, limiting their ability to proliferate and fully colonize the new environment (Theoharides & Dukes, 2007). Together, these results suggest that vegetative traits alone may not fully explain colonization dynamics and highlight the importance of reproductive traits during the establishment phase. The ability to reproduce in a new environment may therefore represent an important component of habitat colonization.

While the role of vegetative traits as enablers of habitat shifts has been increasingly discussed (e.g., Verdú et al., 2003; Ackerly, 2004; Simon et al., 2009; Salvo et al., 2010; Keeley et al., 2011; Zanne et al., 2014; Velasquez-Puentes et al., 2023; Li et al., 2025), less attention has been given to the potential role of reproductive traits in this context (but see, Leishman & Westoby, 1994; Pannell et al., 2015). Reproductive traits are often assumed to be primarily adaptive (Stebbins, 1970; Dellinger, 2026; e.g., Hamilton & Wessinger, 2022; Velasquez-Puentes et al., 2023; Kopper et al., 2024; Dale et al., 2026), but our results suggest that they may also contribute to habitat colonization (e.g. Bonnaudet et al., 2026). In line with this idea, de Frens et al. (2025) found that, across angiosperms, seed dispersal mode can act as an enabler trait for biome shifts. Specifically, they showed that lineages with animal-mediated dispersal more frequently transition into tropical biomes than those with abiotic dispersal, whereas lineages with abiotic dispersal more often shift into arid and temperate biomes than those relying on animal-mediated dispersal.

More broadly, our findings indicate that trait–environment matching can arise from both enabler traits, which may facilitate colonization, and adaptive traits that evolve after habitat transitions. Our results, together with a growing body of empirical evidence (Verdú et al., 2003; Ackerly, 2004; Simon et al., 2009; Salvo et al., 2010; Keeley et al., 2011; Zanne et al., 2014; Velasquez-Puentes et al., 2023; Li et al., 2025; de Frens et al., 2025), suggest that enabler traits may be relatively common and, in some contexts, comparable in frequency to adaptive traits. This highlights the importance of caution when inferring adaptation solely from trait–environment matching without explicitly considering the timing of trait evolution.

### Colonization of harsher environments may require more enabler traits

A key and initially unexpected result was that transitions into harsher environments (X1; characterized by open canopies, lower precipitation and temperature, and greater climatic variability) were more strongly associated with enabler traits, whereas transitions into less harsh environments (X0) showed a relatively greater contribution of adaptive traits. Although somewhat surprising at first, this pattern is consistent with the idea that harsh abiotic conditions impose stronger ecological filters (Pierce et al., 2007; Grime & Pierce, 2012; Tameirão et al., 2021), possibly requiring lineages to possess suitable trait combinations prior to successful colonization (e.g., Ackerly, 2004; Verdú et al., 2003; Salvo et al., 2010; Simon et al., 2009; Keeley et al., 2011; Zanne et al., 2014).

These findings align with the hypothesis proposed by Dale et al. (2024) that trait innovations should be more common during shifts into more extreme biomes. In their study of New Zealand woody flora, the relatively small number of biome transitions associated with trait change occurred primarily during major shifts between forest and alpine environments. However, they did not explicitly test whether trait evolution preceded or followed these transitions. Our results extend this perspective by suggesting that shifts into harsher environments may be more strongly characterized by enabler traits, indicating that pre-existing trait configurations could play an important role in enabling these transitions. Together, these results support an *a posteriori* hypothesis that we encourage future studies to test: colonization of harsh environments may require a greater number of enabler traits, reflecting stronger environmental filtering and tighter constraints on establishment. In contrast, milder environments may allow greater post-colonization trait evolution, resulting in a higher prevalence of adaptive patterns.

## Conclusions, limitations, and future directions

This study has several limitations. First, our analyses are restricted to a single lineage within a tropical region. Although this lineage represents a highly functionally and ecologically diverse system (Oliveira-Filho & Fontes, 2000; Staggemeier et al., 2017; POWO, 2024; Cunha, 2024), the patterns observed here should be generalized with caution, and broader comparative studies across different plant groups and biogeographic contexts will be necessary to assess the generality of our results. Second, the discretization of continuous traits may have reduced the resolution of trait variation and obscured more subtle evolutionary patterns. We encourage further development of generalized dynamic phylogenetic models (GDPMs, Ringen et al., 2026) to allow testing for asymmetric sequences of change in trait evolution contingencies, which may allow future studies to distinguish between enabler and adaptive traits using continuous data. Third, we treated traits as independent variables, whereas interactions among traits may play an important role in shaping evolutionary trajectories (Olson & Miller, 1958; Berg, 1960; Primack, 1987); incorporating trait integration and covariance into future analyses may reveal additional patterns not captured here. The same applies to environmental variables, which were tested separately in our study despite often being correlated. We chose this approach to distinguish subtle differences among climatic gradients that could produce distinct enabler or adaptive patterns. Nevertheless, we encourage future studies to implement multivariate models that incorporate multiple environmental variables simultaneously. Because GDPMs already support multivariate trait evolution, they provide a promising framework for this line of research (Ringen et al., 2026). Finally, we did not directly test evolutionary processes. Instead, our inferences are based on observed patterns interpreted in light of our *a priori* hypotheses. A more explicit distinction between enabler and adaptive processes will require approaches that directly assess the role of traits in facilitating environmental transitions, potentially including experimental studies using plant models with short generation times (e.g., Alzate et al., 2020). Combining macro-and microevolutionary approaches may further improve our understanding of how traits influence habitat transitions.

Overall, our results indicate that habitat shifts in Neotropical Myrtaceae are shaped by a combination of enabler and adaptive patterns, with differences between vegetative and reproductive traits and a strong dependence on environmental context. Vegetative traits tend to function primarily as enablers of habitat shifts, whereas reproductive traits appear to play a dual role, contributing both to establishment and to post-colonization adaptation. The observation that reproductive traits may also act as enablers, contrary to our initial expectations, highlights a potentially underexplored dimension of trait evolution during habitat transitions and suggests new avenues for research. More broadly, these findings point to a greater role of reproductive traits in colonization processes than is typically assumed. In addition, the balance between enabler and adaptive patterns varies across environmental gradients. Transitions into harsher habitats appear to rely more strongly on pre-existing trait configurations, leading us to propose that such conditions may require a greater number of enabler traits for successful colonization. We encourage future studies to explicitly test this hypothesis across multiple clades and environmental contexts.

## Acknowledgments

We thank the staff and curators of the herbaria ALCB, CEPEC, ESA, IAN, MBM, MG, RB, SP, SPF, UB, and UFRN for logistical support. We thank Anselmo Nogueira, Eduardo Koerich Nery and Diogo Borges Provete for their insightful comments on an early version of this manuscript. We are also grateful to Vieira BM for help with figure design. This study was financed in part by the Coordenação de Aperfeiçoamento de Pessoal de Nível Superior - Brasil (CAPES) - Finance Code 001 and by the Instituto Serrapilheira (grant number Serra-R-2111-39858).

## Author contributions

YK, TV, and VGS conceived and designed the study. YK collected the data. YK and JDB performed the analyses. TV and VGS supervised the study, and VGS was responsible for funding acquisition, project administration and resource allocation. YK drafted the initial manuscript, and all authors contributed to the final version of the manuscript.

## Data availability

The data supporting the findings of this study are available in Table S5 at the Supporting Information. Additionally, all data and code are openly accessible at https://github.com/ykilsztajn/habitat_shift_myrtaceae

## Supporting Information

Additional Supporting Information may be found online in the Supporting Information section at the end of the article.

**Fig. S1** PCA biplot showing variation in flower size metrics and trait loadings.

**Fig. S2** Relationship between observed and predicted leaf mass per area (LMA).

**Fig. S3** Distribution of climatic variables across biome types based on species mean values.

**Fig. S4** Frequency of trait states and their effects on transition-index null expectations.

**Table S1** Vegetative and reproductive traits, including their functional significance and expected environmental associations.

**Table S2** Number of herbarium specimens used for trait and occurrence records for each species.

**Table S3** Associations between plant traits and environmental variables.

**Table S4** Observed and null-model transition indices for each trait–environment pair and index direction.

**Table S5** Species mean trait and environmental data used in the analyses.

**Methods S1** Custom Python script for automated extraction of leaf traits using OpenCV.

**Methods S2** Predictive model outputs used to estimate missing LMA values from other leaf morphological traits.

